# Sometimes extracellular recordings fail for good reasons

**DOI:** 10.1101/2025.07.01.662690

**Authors:** Karoline Horgmo Jæger, Aslak Tveito

**Affiliations:** Simula Research Laboratory, Norway

## Abstract

Excitable cells are commonly studied via the extracellular potentials (EPs) they generate, which underlie signals in electroencephalography (EEG), electrocardiography (ECG), and multielectrode array (MEA) recordings. However, some excitable systems produce little or no detectable EPs, for reasons that remain poorly understood. Here we show mathematically that homogeneous excitable cells and tissues – with spatially uniform ion channel distributions and no external stimulation – are extracellularly silent even in the presence of full action potentials. Specifically, an isolated, autonomous cell with uniform membrane properties generates zero EP, independent of shape, kinetics, or model complexity. The result extends to coupled cells provided the tissue remains fully homogeneous. EPs emerge only from spatial inhomogeneities, propagating electrical waves, or applied currents. We demonstrate the physiological relevance of this principle in Purkinje neurons, where clustering of sodium channels enables ephaptic synchronization, while uniform cells remain asynchronous and undetectable extracellularly. We further show that connected human induced pluripotent stem cell–derived cardiomyocytes (hiPSC-CMs) and pancreatic *β*-cells exhibit EPs in proportion to cellular or tissue-level heterogeneity. These findings offer a unifying explanation for the observed silence of some excitable cells and are consistent with experimental reports of strong intracellular signals accompanied by weak or absent EPs.

---

Excitable cells are ubiquitous in physiology, and their electrical fingerprints are heavily investigated to understand the properties of the associated tissue. Of central interest in electrophysiology are the intracellular potential *u_i_*, the extracellular potential *u_e_*, and their difference – the transmembrane potential *v* = *u_i_* −*u_e_*. While *v* is the physiologically most relevant quantity, it is often difficult to measure directly. As a result, *u_e_* plays a central role in clinical investigations of the heart (ECG), the brain (EEG), and in research at the cellular level (MEA).

Extracellular potentials (EPs) form the basis of tools that remain indispensable in clinical practice, offering low-cost, accessible, system-level readouts of electrical activity in the heart and brain. Yet despite their success, EP recordings are not without limitations. Some electrically active tissues appear nearly silent, escaping detection even when robust transmembrane signals are present. For instance, for the *β*-cells of the pancreatic islet, EPs are approximately three orders of magnitude smaller than the transmembrane action potentials (APs) [1], limiting the amount of physiological detail that can be extracted from extracellular recordings [2]. Similar challenges are encountered in MEA-based assays of human induced pluripotent stem cell–derived cardiomyocytes (hiPSC-CMs). For low-density cultures or suboptimal signal filtering, extracellular signals are often small or distorted, reducing the interpretability of the measurements [3]. Furthermore, it has been observed that large populations of neurons remain undetected in extracellular recordings despite appearing metabolically intact [4, 5]. These observations are typically explained in terms of intrinsic silence or sparse firing. Here, we explore a third possibility: that some neurons may be electrically active, generating APs, yet produce no measurable EPs due to a spatially uniform distribution of ion channels. In this view, extracellular silence arises not from a lack of activity, but from biophysical conditions that render the activity undetectable by extracellular means.

The EP plays a particularly important role for Purkinje neurons, which are intrinsic pacemakers that synchronize through ephaptic coupling [6]. This process relies on strong local extracellular fields generated by a high density of sodium channels in the axon initial segment (AIS), where action potentials are initiated [7, 8, 9].

Here, we use mathematical models of excitable cells and tissues to explain the observations described above. For neurons, we apply the classical cable equation coupled to the electrostatic (Laplace) equation [10, 11], as well as the more accurate extracellular (E), membrane (M), intracellular (I) – EMI – model [12, 13]. For cardiac tissue, we use the bidomain model [14] and the Kirchhoff network model (KNM) [15]. We also use the KNM to model a collection of pancreatic *β*-cells.

Previous studies have shown, for a specialized spherical neuron, that EPs vanish when ion channels are uniformly distributed over the membrane [16]. We show that the disappearance of EPs under uniform ion channel distribution is not restricted to this special case. Rather, the result holds in general: if ion channels are uniformly distributed along the membrane of an autonomous cell, and no external stimulation is applied, the EP vanishes – even during a full transmembrane AP. This finding applies across a broad class of membrane models, independent of cell shape or geometry. Numerical simulations further show that non-uniform ion channel distributions or localized stimulation can generate intracellular voltage gradients, which in turn give rise to extracellular signals. We also show that the same principle applies to tissues: uniformity suppresses EPs, while spatial inhomogeneity or localized stimulation reintroduces them. We find that clustering of sodium channels can produce extracellular fields strong enough to synchronize neighboring neurons. These results define concrete, testable physiological hypotheses.

## Neurons modeled by cable theory

We start with the classical *cable equation* of neuroscience, expressed as

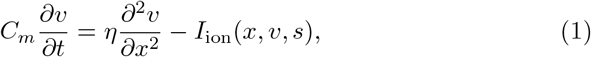

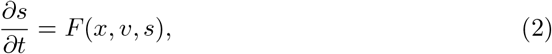

where *t* denotes time, *x* the spatial coordinate, *C_m_* the specific membrane capacitance, and *η* denotes a coefficient derived from the intracellular conductivity and cell geometry [12]. The function *I*_ion_(*x, v, s*) represents the total ionic current density, and *F* (*x, v, s*) governs the evolution of the state variables *s*, which describe ion concentrations and gating variables. For further background and derivations of the cable equation, see [17, 18, 11].

We consider a neuron with an intracellular volume Ω*_i_*, embedded in an extracellular space Ω*_e_*, and separated by the cell membrane Γ. Based on the solution of the cable equation, the EP can be computed (see, e.g., [19, 10, 12]) by solving the electrostatic equation

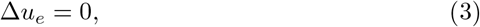

with the boundary condition

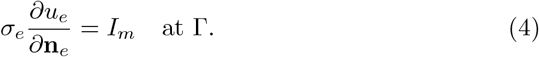

Here, *σ_e_* is the extracellular conductivity, **n***_e_* is the outward normal vector from the membrane into the extracellular space, and *I_m_* = *η∂*^2^*v/∂x*^2^ is the membrane current density. We assume that *u_e_* vanishes at the outer boundary of Ω*_e_*.

Suppose the cable is cylindrical and completely uniform. Then the functions *I*_ion_(*x, v, s*) and *F* (*x, v, s*) reduce to 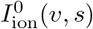 and *F* ^0^(*v, s*), respectively. Consider the following system of ordinary differential equations (ODEs)

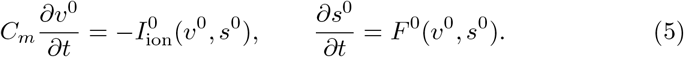

Since the cable is uniform, the functions *v*^0^(*t*) and *s*^0^(*t*) also solve the full cable system (1) and (2). Because all spatial derivatives vanish, the membrane current *I_m_* is zero, and then it follows from the electrostatic equations (3) and (4) that the extracellular potential *u_e_* is identically zero throughout the domain Ω*_e_*. We conclude that in this uniform case, the triplet (*v, s, u_e_*) = (*v*^0^*, s*^0^, 0) solves the system (1)–(4).

If *I*_ion_ and *F* are chosen such that the solution *v*^0^*, s*^0^ represents a pacemaker exhibiting full-scale APs (e.g., with *v*^0^ ranging from 80 mV to +60 mV), the cell generates a complete AP while producing no extracellular signal.

A neuron with uniform membrane dynamics thus produces no EP and is invisible to extracellular measurement devices. This changes markedly in the presence of inhomogeneity, as illustrated in Figure 1, where increasing the spatial concentration of sodium channels in localized regions amplifies the EP. Furthermore, if spatially localized stimulation is applied to the neuron – e.g., in the form of synaptic input – it introduces spatial inhomogeneities even in the case of a uniform distribution of ion channels, thereby generating an EP (see rightmost panel of Figure 1).

**Figure 1:**
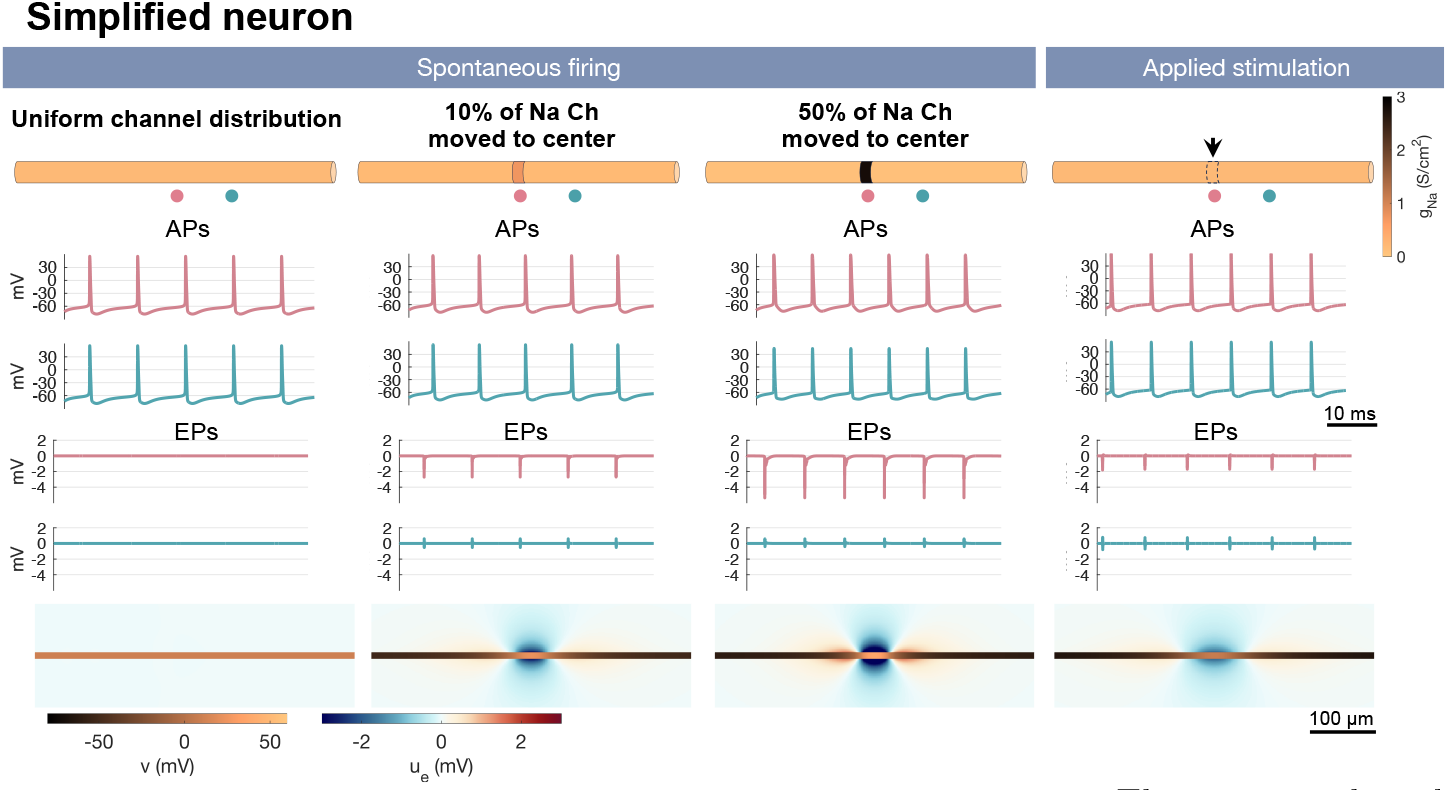
Cable equation computations of a simplified neuron. The neuron is shaped as a cylinder with diameter 10 *µ*m and length 1000 *µ*m. In the left column there is a uniform distribution of ion channels along the cell membrane. Spontaneously firing action potentials (APs) are reported for two points along the cell. Below, the extracellular potential (EP) is shown and is zero. The EP is computed using the point source method (see the Methods section or, e.g., [11, 12]). In the lower panel, a snapshot of the transmembrane potential and EP are shown for a point in time when the cell fires. Here, we focus on the center 500 *µ*m of the neuron. In the second column, mild concentration of sodium channels (Na Ch) in the center 40 *µ*m of the neuron results in a small but non-zero EP. In the third column, further concentration of sodium channels increases the amplitude of the EP. In the fourth column, applied stimulation in the center of a neuron with a uniform distribution of ion channels yields a non-zero EP. The stimulation is applied with a 120 Hz frequency.

Applied stimulation

## Cardiac tissue modeled by the bidomain equations

The *bidomain model* plays a foundational role in computational cardiac electrophysiology, much like the cable equation does in neuroscience. It is widely regarded as the gold standard for simulating electrical activity in cardiac tissue. A recent review highlights its growing relevance for clinical applications [20]. We adopt the standard formulation of the bidomain model:

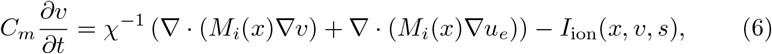

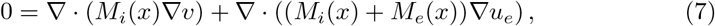

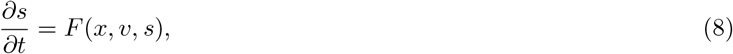

see, e.g., [14, 21]. The system is posed on a spatial domain Ω with boundary *∂*Ω. Here, *M_i_*(*x*) and *M_e_*(*x*) denote the intracellular and extracellular bidomain conductivities, and *χ* is the surface-to-volume ratio of the cell membrane. We impose no-flux boundary conditions for *v*, and a Dirichlet condition *u_e_* = 0 mV on *∂*Ω. Note that all quantities are defined throughout the entire domain Ω, since the bidomain model is derived via homogenization of the underlying discrete cell structure; see [14].

The variables *v* and *s*, and the functions *I*_ion_ and *F*, are defined in the same manner as for the cable model above. However, the underlying biophysics is quite different: while the formal structure of the equations is preserved, cardiac models typically involve longer-duration APs with distinct plateau phases and calcium cycling. Consequently, *I*_ion_ and *F* describe different sets of ionic currents and gating variables than in neuronal models.

Let *v*^0^(*t*) and *s*^0^(*t*) be the solution of the ODE system (5), corresponding to a homogeneous (spatially uniform) excitable cell model. Then, by direct substitution, the triplet (*v, s, u_e_*) := (*v*^0^*, s*^0^, 0) satisfies the full bidomain system (6)–(8) in the spatially homogeneous case. This construction corresponds to a completely homogeneous and autonomous tissue that exhibits full transmembrane dynamics without generating any EP. In this case, the bidomain model yields the same qualitative result as in the single-cell cable model above: the tissue produces full-scale APs, yet remains electrically silent when measured from the extracellular space.

In Figure 2, we show simulations of the electrophysiology of a sinoatrial node tissue sample (cf. [22]). The membrane dynamics are taken from [23], capturing the intrinsic pacemaker activity of sinoatrial cells. In the left panel, a completely homogeneous tissue continues to exhibit autonomous pacemaker activity, but no detectable EP is generated. When moderate inhomogeneities in membrane properties are introduced in four distinct regions, a non-zero EP emerges. This effect is further amplified in the right panel, where the degree of inhomogeneity is increased.

**Figure 2:**
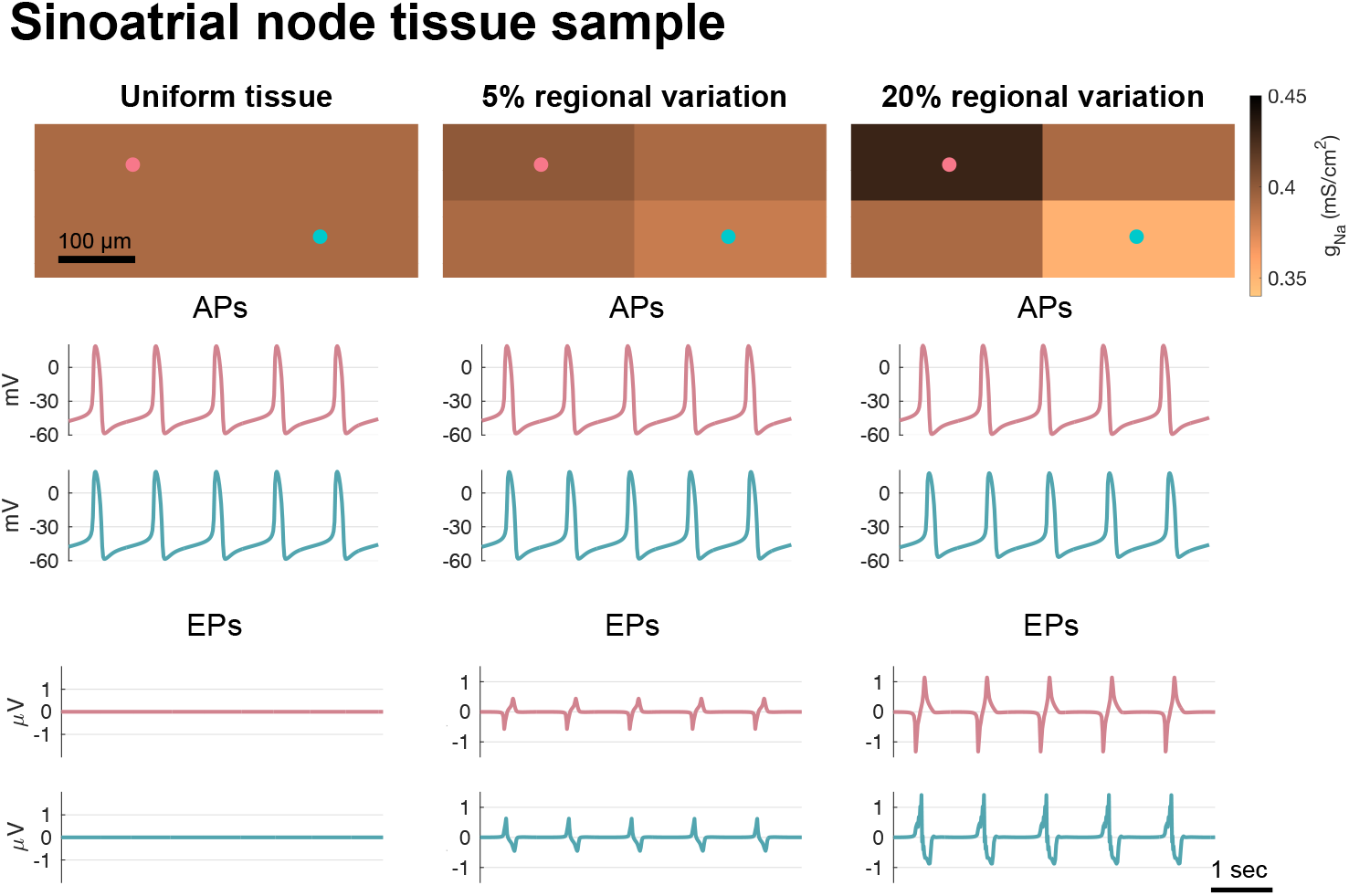
Bidomain model simulations of a sinoatrial node (SAN) tissue sample. A small cube of SAN tissue is modeled using the bidomain model equations. In the left column, the tissue is completely homogeneous, with identical membrane properties in all regions. The transmembrane potential exhibits spontaneous pacemaker activity, but the extracellular potential (EP) remains undetectable. In the middle column, moderate heterogeneity is introduced by varying membrane properties across four spatial regions, leading to a small but measurable extracellular signal. In the right column, larger inhomogeneities result in stronger EPs. The action potentials (APs) and EPs are plotted for the two points indicated by circles in the upper panel. Note that *all* major ion channel conductances are varied between regions like illustrated for *g*_Na_ in the upper panel (see the Methods section).

## Heart-on-a-chip systems modeled by the Kirchhoff Network Model (KNM)

Human induced pluripotent stem cell–derived cardiomyocytes (hiPSC-CMs) have become an important tool for studying human cardiac electrophysiology [24, 25, 26, 27], including the effects of pharmaceutical compounds [28, 29, 30]. In such applications, drug action is often inferred by observing changes in biomarkers and performing model-based inversion [31, 32, 33]. When EPs are used in this context, the bidomain model has been a standard modeling framework [34, 35, 36, 33, 37]. However, this model represents tissue as a homogenized medium and does not resolve individual cells explicitly [38].

To address these limitations, more detailed models have been developed. The extracellular–membrane–intracellular (EMI) model represents the spatial domains of each cell explicitly and offers micrometer-level resolution [39]. Due to its computational cost, alternative cell-based approaches have also been proposed, including the Kirchhoff Network Model (KNM), which treats each cell as a computational unit while still capturing key spatial effects [15, 40, 41]. In a recent benchmark, the KNM completed a typical simulation in 7 seconds, compared to 28 minutes for the bidomain model and 16 hours for the EMI model [42].

We now consider the KNM, in which each cell *k* is represented by a single compartment with transmembrane potential *v^k^*, and an associated extracellular compartment with extracellular potential 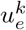. The model equations, derived from Kirchhoff’s current law [15, 40], are:

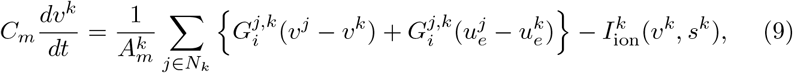

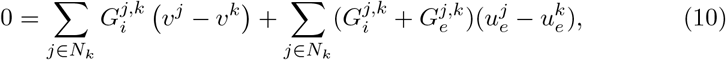

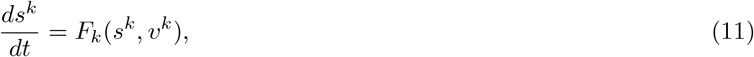

where *N_k_* denotes the neighbors electrically connected to cell *k*, and 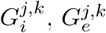 represent intracellular and extracellular conductances. The parameter 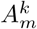 denotes the membrane area of cell *k*. The membrane dynamics represented by the functions *I*_ion_ and *F* are based on [43] (the immature case). The KNM equations capture the evolution of transmembrane potential, the extracellular potential, and subcellular dynamics in a simplified but spatially explicit network of cells.

Motivated by the examples above, we define (*v*^0^*, s*^0^) to be the solution of the following system of ODEs

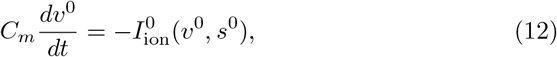

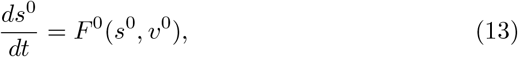

and assume that all cells are identical. Then, assuming homogeneous boundary conditions as above, the triplet 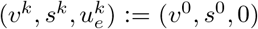 satisfies the full KNM system (9)–(11) for all cells *k*. This demonstrates that a fully homogeneous, autonomous tissue modeled with the KNM framework produces no EP – again confirming the main result: electrical activity can occur without producing any extracellular signature.

In Figure 3, we show results of KNM simulations of a heart-on-a-chip system with different degrees of variation in cell properties. We observe that for no cell variation, no EP is produced despite robust AP firing. Furthermore, small variations in cell properties give rise to small EPs, whereas large cell-to-cell variations or localized stimulation produce strong EPs.

**Figure 3:**
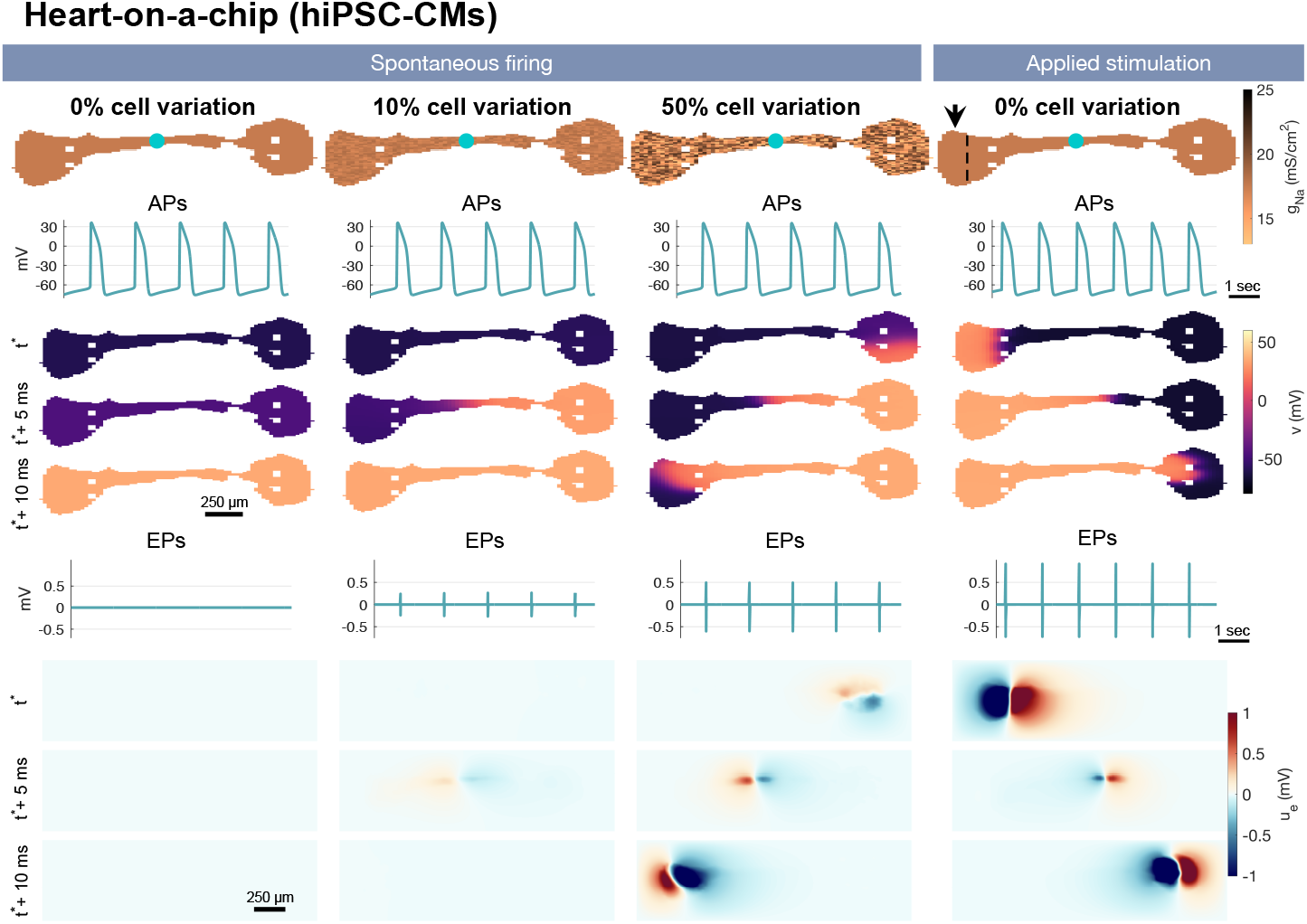
KNM simulation of a heart-on-a-chip. The chip consists of hiPSC-CMs connected by gap junctions. The chip geometry, parameters and membrane dynamics model are adapted from [41] and described in the Methods section. In the left panel, there is no variation between the cells and spontaneous action potentials (APs) are fired uniformly across the chip. The AP and extracellular potential (EP) traces are taken from a point in the center of the cell collection (marked by a blue circle in the upper panel). In addition, we show snapshots of the transmembrane potential and the EP at three points in time close to AP firing. For a chip with uniform cell properties, the EP is zero. In the second panel, 10% cell-to-cell variation is introduced by altering the conductance of all transmembrane currents. This gives rise to an excitation wave and weak EPs. In the third panel, the variation is increased to 50%, producing a stronger propagating wave accompanied by more pronounced EPs. In the rightmost panel, a 1 Hz stimulation current is applied to the left side of the chip, initiating a robust excitation wave and strong EPs throughout the tissue.

Recording EPs from hiPSC-CMs in culture wells is challenging due to low signal amplitudes, immature electrophysiological properties, and variability in how the cells are organized. Poor electrode contact, unclear waveforms, and the need to discard weak signals after recording make interpretation difficult [3, 44]. However, as we show here, even fully functional and synchronized hiPSC-CMs may produce little or no extracellular signal if the cells have equal membrane properties. This suggests that some measurement problems may reflect real electrical invisibility rather than technical failure.

## Pancreatic *β*-cells simulated using KNM

Pancreatic *β*-cells are electrically excitable endocrine cells that secrete insulin in response to elevated blood glucose. Their electrical activity typically takes the form of bursts of APs, tightly coupled to intracellular calcium oscillations and metabolic feedback mechanisms [45, 2]. While patch-clamp recordings have been the standard tool for probing their electrophysiology, EP recordings have also been used, especially in MEA configurations, to monitor collective electrical activity *in vitro* [1].

To investigate the generation of EPs in these cells, we use the KNM structurally identical to the one used for hiPSC-CMs above, but parameterized for human *β*-cell electrophysiology using the membrane model taken from [46]. A key feature of this model is the inclusion of calcium-activated potassium channels, which regulate spiking and bursting activity by responding to submembrane calcium levels. In addition, slow bursting is driven by glycolytic oscillations, introduced as a separate metabolic feedback loop. We consider a three dimensional collection of 1000 such *β*-cells where neighborings cells are connected by gap junctions (see Figure 4).

**Figure 4:**
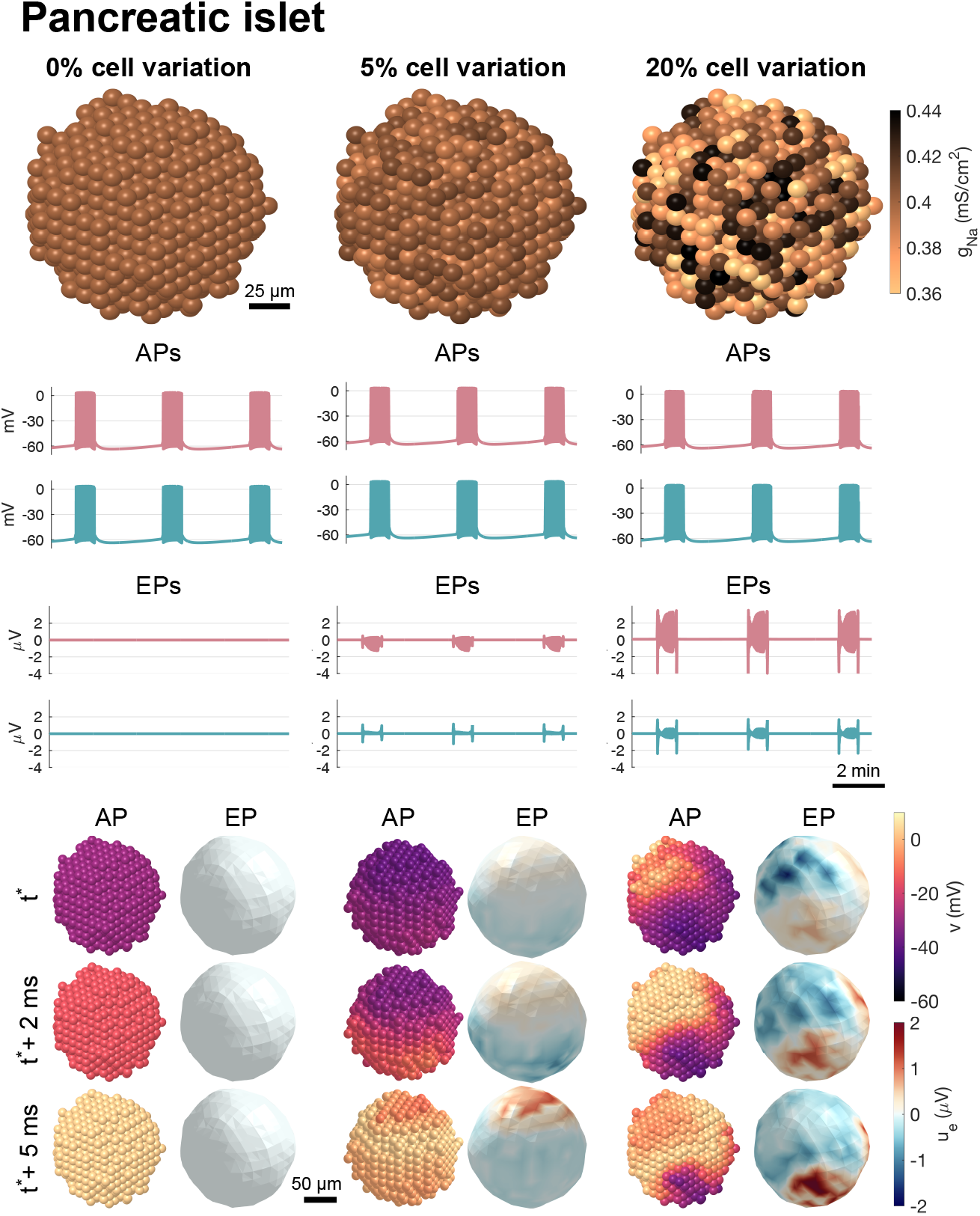
KNM simulation of a pancreatic islet consisting of *β*-cells. The membrane dynamics of the *β*-cells are modeled using the Riz et al. model [46] and the cells are connected to neighbours by gap junctions. In the left panel, there is no variation between the cells. Spontaneous action potentials (APs) are fired in bursting periods, but no extracellular potentials (EPs) are generated. The red lines show the solutions for the cell with the largest EP amplitude, and the blue lines show the solutions for the cell with the smallest EP amplitude. When no EP is produced, this corresponds to two arbitrary cells. In the lower panels, the membrane potential and the EP on the surface of the islet are depicted at three points in time during a single AP firing. In the left panel, all cells are equal, the transmembrane potential is uniform and the EP is zero. In the middle panel, cell variation is introduced by varying the value of *g*_Na_, *g*_CaL_, and *g*_KATP_ between each cell. Again, APs are fired, and now weak EPs are visible. In the lower panel, small transmembrane potential gradients are visible during bursting. In the right panel, more extensive variations are introduced between cells, resulting in more prominent spatial transmembrane potential gradients and stronger EPs. The model set-up used in these simulations is described in detail in the Methods section.

As in the heart-on-a-chip case, it follows directly that homogeneous intracellular and extracellular conductivities yields zero EP. In Figure 4, numerical simulations illustrate the relationship between spatial inhomogeneity and the resulting EPs. When the density of active membrane conductances varies across the cell cluster, propagating depolarizations and EPs occur and the strength of the EP increases as the degree of inhomogeneity increases.

## Cerebellar Purkinje neurons simulated using the EMI model

Cerebellar Purkinje neurons are large, autonomous neurons that play a central role in motor coordination. Their spontaneous firing and ability to synchronize with neighboring cells via ephaptic coupling are critically dependent on a high concentration of voltage-gated sodium channels in the axon initial segment (AIS) [7, 8]. These neurons thus provide a compelling physiological case where localized ion channel clustering governs extracellular signal strength, and thereby cell-to-cell interaction via the extracellular space.

The standard framework for understanding the electrodynamics of neurons is the cable equation, (1), where the EP is computed in a separate post-processing step using the electrostatic equation, (3). However, this decoupling neglects the dynamic feedback between the intracellular, membrane and extracellular domains and is therefore only an approximation [12]. A more accurate approach is offered by the EMI model, which resolves the full electrodynamics by directly coupling the extracellular (E), membrane (M), and intracellular (I) compartments.

For a single cell, the EMI model reads:

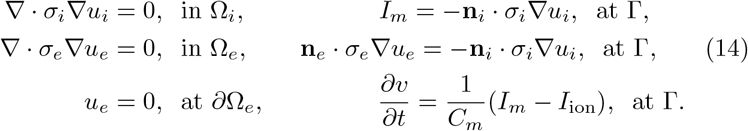

see, e.g., [47, 48, 12]. Here, *u_i_, u_e_*, and *v* = *u_i_* −*u_e_* denote the intracellular, extracellular, and membrane potentials, defined in the domains Ω*_i_*, Ω*_e_*, and on the membrane surface Γ, respectively. The outer boundary of the extracellular space is denoted by *∂*Ω*_e_*, and **n***_i_* and **n***_e_* are the outward-pointing unit normal vectors of Ω*_i_* and Ω*_e_*. As above, the parameters *σ_i_* and *σ_e_* represent the intracellular and extracellular conductivities, and *C_m_* is the specific membrane capacitance. The total ionic current density is given by the function

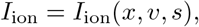

where *x* is the three-dimensional spatial coordinate, and *s* denotes ion concentrations and gating variables governed by

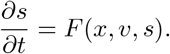

We now consider the autonomous case where the ion channels are uniformly distributed along the cell membrane, and all spatial gradients vanish, with *v*^0^(*t*) and *s*^0^(*t*) solving the membrane ODE system (12), (13). By substituting *v*^0^ and *s*^0^ into the EMI system, we find that the triplet (*v, s, u_e_*) = (*v*^0^*, s*^0^, 0) satisfies the full equations in the case of spatially uniform channel distributions and no external stimulation. The cell is, in this case, extracellularly invisible: its activity produces no EP. Notably, this result does not rely on any specific assumptions about the three-dimensional geometry of the cell.

This theoretical result is illustrated by EMI-based simulations of a morphologically detailed Purkinje neuron model (see Figure 5). When ion channels are non-uniformly distributed across the cell (e.g., with a high density of sodium channels in the AIS), the cell produces spontaneous APs and EPs, but when the ion channels are uniformly distributed across the cell, no EPs are generated despite spontaneous AP firing.

**Figure 5:**
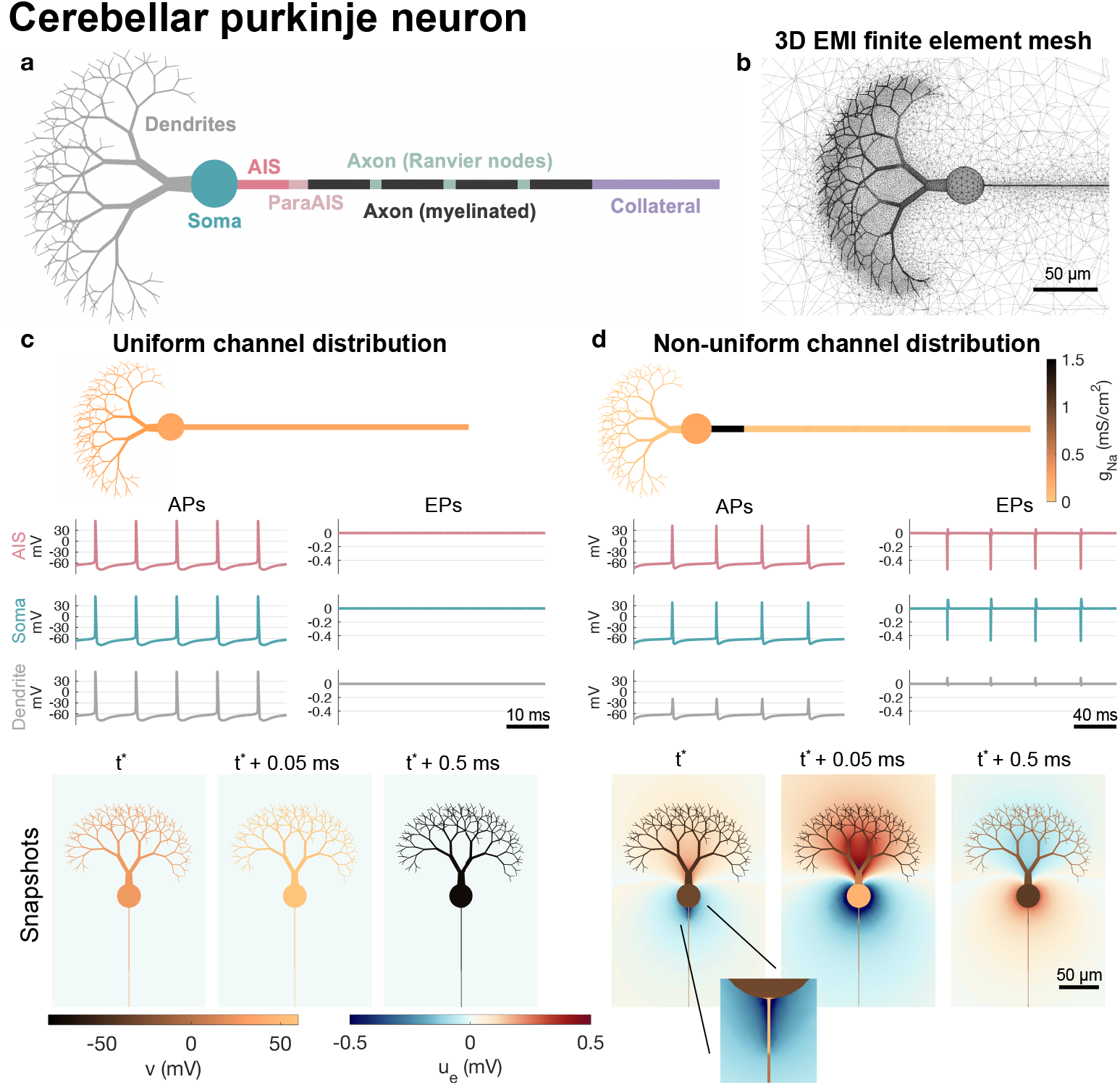
EMI model simulations of a cerebellar purkinje neuron. **a.** Illustration of the different geometrical parts of the neuron. The model parameterizations for each neuron section is based on [7] and specified in the Methods section. **b.** Illustration of the three dimensional finite element mesh close to the soma and dendrites used in the EMI model simulations. **c.** Simulation results for a uniform channel distribution. The channel densities of the soma are applied in the entire neuron. The average membrane potential (APs) and extracellular potential (EPs) over different parts of the neuron is illustrated. More specifically, we consider the axon initial segment (AIS), the soma, and the outermost part of the dendrites (diameter *<* 3.5 *µ*m). Even though APs are clearly present, no EPs are visible. In the lower panel, snapshots of the transmembrane potential and EP is illustrated at the point in time when the AIS membrane potential reaches a value above 0 mV, *t*\*, and 0.05 ms and 0.5 ms after *t*\*. Again, no EP is visible. **d.** Same setup as in panel c, but for a physiological, nonuniform distribution of ion channels, based on [7]. In this case, the neuron displays both APs and EPs.

## Ephaptic synchronization requires spatial inhomogeneity

The preceding sections demonstrate that uniform excitable cells generate no EP, and hence remain electrically invisible to their surroundings. This has direct implications for ephaptic coupling, where cells influence each other via the extracellular field. In the absence of spatial inhomogeneities – such as localized ion channel clustering – no ephaptic interaction is possible. We illustrate this principle using cerebellar Purkinje neurons, where ephaptic synchronization is observed only when sodium channels are concentrated in the AIS.

For cardiomyocytes, the role – and even the existence – of ephaptic coupling between neighboring cells has long been debated; see, e.g., [49, 50]. A similar debate exists for neurons, though shaped by different physiological considerations. Nonetheless, experimental evidence confirms that cerebellar Purkinje neurons can synchronize via ephaptic coupling – that is, through interactions mediated by the EP [6]. This implies that extracellular fields are not merely observational artifacts, but can play a direct physiological role.

In Figure 6, we show that two Purkinje neurons can synchronize via the EP. When sodium channels are uniformly distributed (left panels), no extracellular field is generated and the APs remain asynchronous. In contrast, when sodium channels are clustered in the AIS – as observed experimentally – synchronization emerges rapidly via the extracellular field (right panels). This finding is consistent with experimental results showing that sodium channel clustering in the AIS is necessary to generate strong extracellular fields and support synchronization [9, 6].

**Figure 6:**
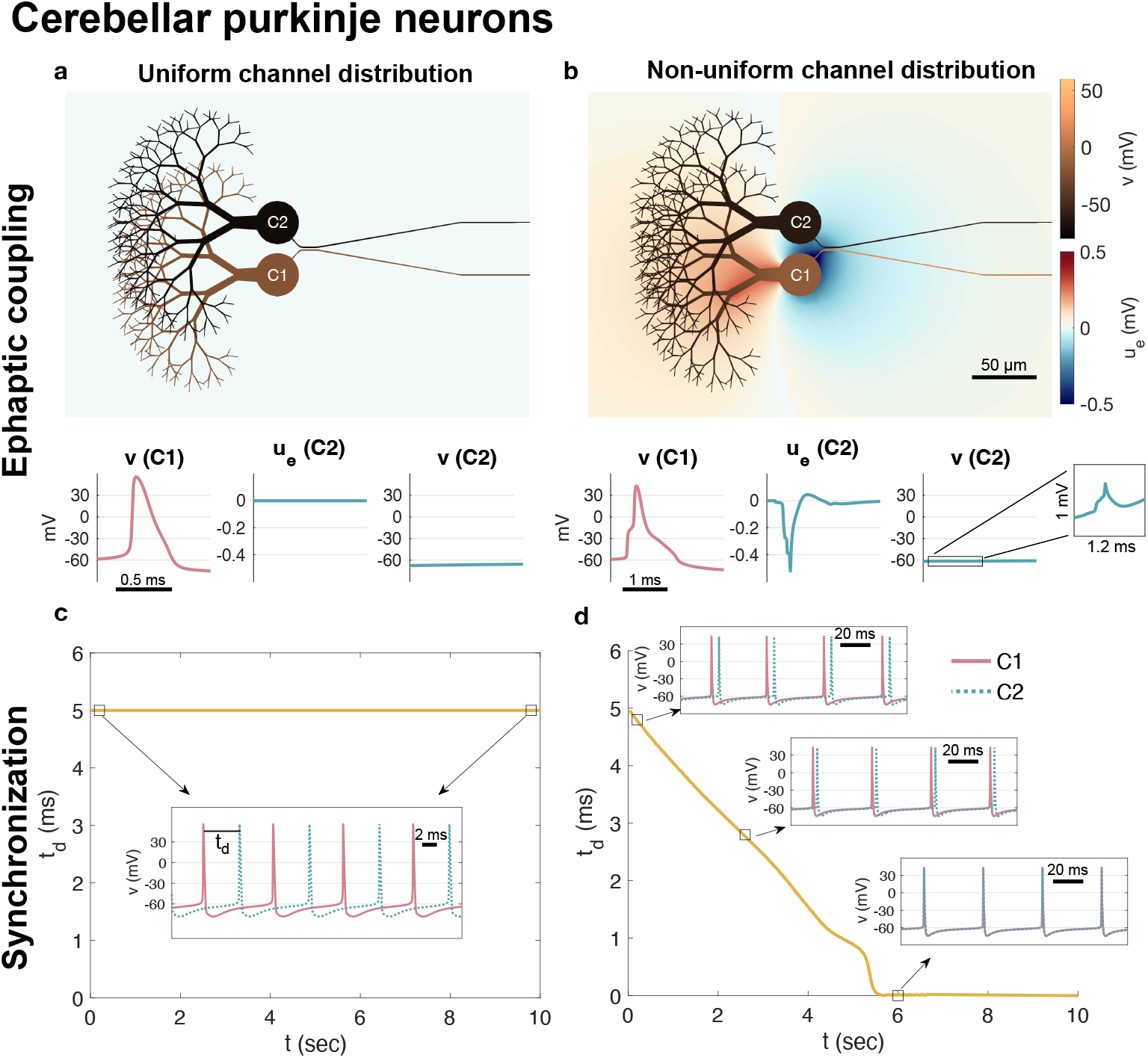
Synchronization in EMI model simulations of two cerebellar purkinje neurons. The neuron geometry and parameterizations are the same as in Figure 5, but the neurons are set up such that the lower neuron, C1, fires action potentials (APs) 5 ms before the upper neuron, C2. **a.** Both neurons have a uniform ion channel distribution. A solution snapshot is shown at a point in time when C1 fires, and no extracellular potential (EP) is visible. In addition, the average transmembrane potential of the AIS of C1, the average EP of the AIS of C2 and the average transmembrane potential of the AIS of C2 are plotted as functions of time during the time of a C1 AP. **b.** Both neurons have a non-uniform distribution of ion channels based on [7] like in Figure 5d. When C1 fires an AP, a negative EP reaches from the AIS of C1 to the AIS of C2. Below, we observe that this results in a negative EP for C2 and a small positive notch in the transmembrane potential of C2. **c.** As the simulation of the two uniform cells is continued for 10 seconds, the distance between the AP peaks remain constant at 5 ms. **d.** For the non-uniform purkinje neurons, the EP affects the transmembrane potential of the neighboring cell and a gradual synchronization (i.e., a reduction in the AP peak distance) is observed. After 6 seconds of simulation, the two neurons fires synchronously.

## Biophysical origin of extracellular silence

In the mathematical arguments given above, we consistently start by considering a uniform, unstimulated cell with no intracellular spatial gradients. We then deduce, using the cable equation (1)–(2), the bidomain model (6)–(8), the KNM (9)–(11), and the EMI model (14), that the EP remains zero. All these models are derived from Kirchhoff’s current law, which states that the total current into a compartment must equal the total current out. In fact, the main result – that no EP is generated – follows directly from this law and does not rely on any particular partial differential equation.

Figure 7 shows an intracellular compartment Ω*_i_* with left and right axial currents *I_i,l_* and *I_i,r_*, and a membrane current *I_m_*, which includes both the capacitive component 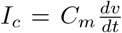 and the ion channel component *I*_ion_. If the intracellular potential is spatially constant, the ohmic axial currents vanish, so *I_i,l_* = *I_i,r_* = 0, and Kirchhoff’s law implies that *I_m_* = 0. With no current *I_m_* flowing across the membrane, the extracellular space, Ω*_e_*, is unaffected so the EP remains zero.

**Figure 7:**
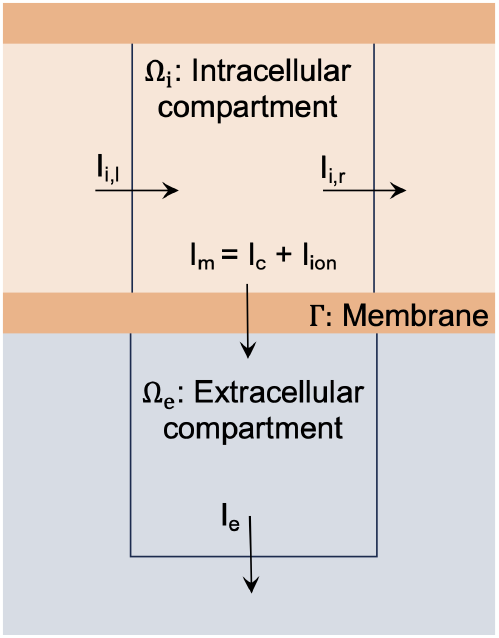
Schematic of intracellular and extracellular compartments used to illustrate Kirchhoff’s current law. In the absence of intracellular voltage gradients, axial currents *I_i,l_* and *I_i,r_* vanish, implying *I_m_* = 0 and therefore no extracellular current *I_e_*. The total membrane current *I_m_* is the sum of the capacitive and ionic components, *I_m_* = *I_c_* + *I*_ion_.

Furthermore, the condition *I_m_* = 0 implies that the capacitive and ionic currents must balance each other: *I_c_* + *I*_ion_ = 0. This gives rise to a system of ODEs, (5), that governs the time evolution of the membrane potential and state variables. In short, a uniform, autonomous excitable cell may exhibit full-scale APs internally while producing no extracellular signal.

## Consistency with experimental observations

As we have seen, a uniform distribution of ion channels along the cell membrane leads, in mathematical models, to an EP that is exactly zero. Physically, however, ion channels are discrete molecular structures, and perfect homogeneity is not achievable. As a result, EPs in real physical systems are expected to remain extremely small – but not strictly zero. Similarly, in computational models, we have shown that EPs computed under the assumption of uniform channel distribution are minuscule, though these are not necessarily strictly identical to zero either, due to round-off and discretization errors. This result holds both for isolated cells with nearly uniform channel distributions and for multicellular assemblies such as cultured cardiomyocytes or pancreatic *β*-cells, which share similar membrane properties and are electrically coupled via gap junctions. In such systems, intracellular signals may be robust, while extracellular recordings show little or no detectable activity. However, our simulations show that when externally stimulated, even homogeneous tissues generate propagating excitation waves that produce significant EPs.

Several studies report cases where EPs are small or absent despite the presence of clear intracellular signals. In [1], microelectrode array (MEA) recordings from intact pancreatic islets revealed extracellular signals on the order of a few microvolts, even during glucose-induced bursting, while corresponding intracellular recordings showed APs of tens of millivolts. Similarly, [51] highlights that although MEAs provide noninvasive and high-throughput recordings, they offer limited information about the shape of electrical signals and cannot resolve subtle changes. In the context of cultured cardiac tissues, [3] demonstrates that field potential amplitude and waveform depend strongly on cell plating density and filtering parameters. In low-density or unstimulated conditions, the field potentials often appear weak or distorted – even when intracellular activity is robust. Notably, when a biphasic electrical stimulus was applied, the resulting extracellular signals became significantly stronger and better aligned with AP dynamics. This observation is consistent with our theoretical result that extracellular signals emerge from stimulation-induced inhomogeneity, even in otherwise uniform systems.

By contrast, other studies show that strong EPs can be recorded from cells with marked inhomogeneities in membrane properties. In [52], vertical plasmonic nanoelectrodes combined with optoporation enabled high-fidelity recording of both intracellular and extracellular signals from cardiomyocytes and neurons. In their experiments, spontaneously firing hippocampal neurons exhibited robust extracellular signals. Viewed in light of our results, this suggests non-uniform ion channel distributions. This interpretation aligns with direct experimental evidence of sodium channel clustering in these neurons: as shown in [53], the AIS of hippocampal pyramidal cells contains a dense accumulation of low-threshold Nav1.6 channels at the distal AIS, which determines the site of AP initiation.

Taken together, these findings suggest that, in autonomous cells that are not externally stimulated, the presence or absence of observable EPs depends not only on the recording technique or signal processing, but on the spatial distribution of cell properties. Our results provide a theoretical explanation for how electrically active cells can remain invisible to extracellular recording methods when ion channels are uniformly distributed. This may help clarify longstanding observations of electrophysiologically silent, yet metabolically active, neurons – sometimes referred to as “silent neurons” or “dark matter” of the brain [4, 5]. While previous work emphasized sparse firing or extreme selectivity, our findings suggest that some of this invisibility may instead reflect uniformly distributed electrical activity that fails to generate EPs. In this sense, “neural dark matter” may not be silent at all – just uniformly active in a way that renders it invisible to extracellular probes.

## Methods

### Cable equation representation of a simplified neuron

In the cable equation representation of a simplified neuron in Figure 1, the neuron is shaped as a cylinder of length 1000 *µ*m and diameter *d* = 10 *µ*m. The transmembrane currents (*I*_ion_ and *F*) are modeled using a slightly altered version of the cerebellar purkinje neuron model from [7]. The alterations from [7] are described below in the specification of the EMI model of a purkinje neuron. We use the transmembrane conductances corresponding to the soma domain (see Table 1). In the case of applied stimulation, a 0.1 ms long constant stimulation current of 3 mA/cm^2^ is applied at 120 Hz frequency in the center 40 *µ*m of the neuron. The specific membrane capacitance is set to *C_m_* = 1 *µ*F/cm^2^ [54]. The parameter *η* is given by

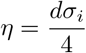

(see, e.g., Chapter 9 in [21]), where *d* is the neuron diameter and *σ_i_* = 8.2 mS/cm [7] is the intracellular conductivity. The extracellular conductivity is set to *σ_e_* = 3 mS/cm^2^ [55], and the extracellular potential is computed using the point source formula

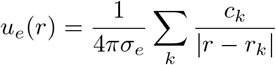

(see, e.g., [11, 55, 12]). Here, *c_k_* is the current source from each compartment, *k*, in the discretization of the cable equation and *|r* −*r_k_|* is the distance between the point where *u_e_* is measured and the center of compartment *k*.

**Table 1.** Overview of lengths, *l*, (in *µ*m) diameters, *d*, (in *µ*m) and conductances of different types of ion channels (in S/cm^2^) in the spatial regions of the purkinje neuron, based on [7]. There are four Ranvier nodes (RN). Between these, there are myelinated regions of length *l* = 100 *µ*m and diameter *d* = 0.7 *µ*m. For the lengths and diameters of the dendric regions, see Table 2.

| | Dendrite<br>($d < 3.5 \mu\text{m}$) | Dendrite<br>($d > 3.5 \mu\text{m}$) | Dendrite<br>($d > 8 \mu\text{m}$) | Soma | AIS | ParaAIS | RN | Collateral |
| --- | --- | --- | --- | --- | --- | --- | --- | --- |
| $l$ | | | | 30 | 17 | 4 | 4 | 100 |
| $d$ | | | | 30 | 1 | 1 | 0.7 | 0.6 |
| Nav1.6 | 0 | 0 | 0.016 | 0.214 | 1.5 | 0 | 0.03 | 0.03 |
| Kv1.1 | 0.036 | 0.0012 | 0.0012 | 0.002 | 0 | 0.01 | 0 | 0 |
| Kv1.5 | 0.0039 | 0.00013 | 0.00013 | 0 | 0 | 0 | 0 | 0 |
| Kv3.3 | 0.3 | 0.01 | 0.01 | 0 | 0 | 0 | 0 | 0 |
| Kv3.4 | 0 | 0 | 0 | 0.05 | 0.01 | 0 | 0.01 | 0.02 |
| Kv4.3 | 0.03 | 0.001 | 0.001 | 0 | 0 | 0 | 0 | 0 |
| Kir2.x | 0 | 0.00001 | 0.00001 | 0.00003 | 0 | 0 | 0 | 0 |
| Kca1.1 | 1.05 | 0.035 | 0.035 | 0.01 | 0 | 0 | 0 | 0 |
| Kca2.2 | 0 | 0.001 | 0.001 | 0.001 | 0 | 0 | 0 | 0 |
| Kca3.1 | 0 | 0.002 | 0.002 | 0.01 | 0 | 0 | 0 | 0 |
| Cav2.1 | 0.001 | 0.001 | 0.001 | 0.00022 | 0.00022 | 0 | 0.00022 | 0.00022 |
| Cav3.1 | 0 | $5 \cdot 10^{-6}$ | $5 \cdot 10^{-6}$ | $7 \cdot 10^{-6}$ | $1 \cdot 10^{-5}$ | 0 | $1 \cdot 10^{-5}$ | $1 \cdot 10^{-5}$ |
| Cav3.2 | 0 | 0.0012 | 0.0012 | 0.0008 | 0 | 0 | 0 | 0 |
| HCN1 | $4 \cdot 10^{-6}$ | $4 \cdot 10^{-6}$ | $4 \cdot 10^{-6}$ | 0.0004 | 0 | 0 | 0 | 0 |

#### Introducing inhomogeneities

We introduce a non-uniform ion channel distribution along the cell by moving a certain fraction, *p* = 0.1 or *p* = 0.5, of the sodium channels to the center 40 *µ*m of the neuron. For the remaining part of the neuron membrane, the sodium channel conductance is set to

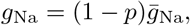

where 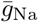 is the default value. In the center 40 *µ*m of the neuron, the sodium channel conductance is set to

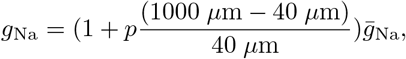

keeping the integral of *g*_Na_ constant for different values of *p* (see, e.g., [56]).

#### Numerical methods

In the cable equation representation of a simplified neuron, the neuron is discretized into compartments of length 1 *µ*m along the longitudinal neuron axis.

The cable equation can then be represented as a coupled system of ordinary differential equations (see, e.g., [12]). This system is solved using the ode15s solver in MATLAB, which uses adaptive time stepping.

### Bidomain model representation of a sinoatrial node tissue sample

In the bidomain model representation of a sinoatrial node tissue sample (Figure 2), we consider a tissue of size 0.5 *×* mm 0.2 mm *×* 0.1 mm, surrounded by an extracellular bath of width 0.1 mm in all spatial directions. We apply no-flow Neumann boundary conditions for the membrane potential and *u_e_* = 0 mV Dirichlet boundary conditions for the EP.

The membrane dynamics (*I*_ion_ and *F*) are modeled using the human sinoatrial node model from [23]. The bidomain model parameters *χ, M_i_* and *M_e_* are set up based on tissue properties, following [57, 40]. That is, we define

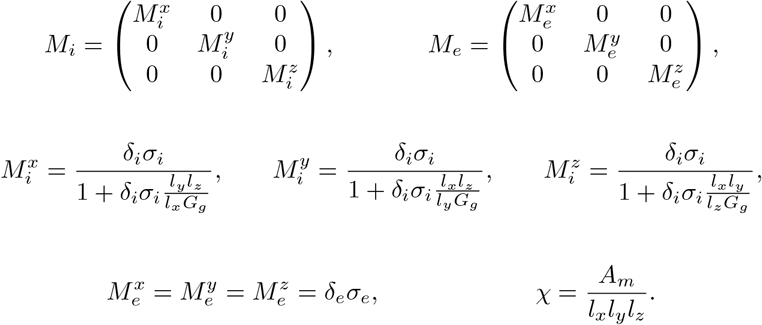

Here, *σ_i_* = 8.2 mS/cm [7] is the intracellular conductivity, *σ_e_* = 15 mS/cm [58] is the extracellular conductivity, and *G_g_* = 1 nS = 10^−6^ mS [59] is the gap junction conductance between to neighboring cells. Furthermore, *A_m_* is the surface area of a cell, *l_x_, l_y_, l_z_* are the lengths of a cell in each spatial direction, and *δ_i_* and *δ_e_* = 1− *δ_i_* are the intracellular and extracellular volume fractions, respectively. From [23], each cell is assumed to be a cylinder of length *l_x_* = 67 *µ*m and diameter *d* = *l_y_* = *l_x_* = 7.8 *µ*m. Assuming that the extracellular space covers the remaining volume *l_x_ × l_z_ × l_z_*, the intracellular volume fraction is 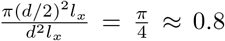. From [23], the capacitance of a single cell is 57 pF, and assuming a specific membrane capacitance of *C_m_* = 1 *µ*F/cm^2^ [54], the membrane area is *A_m_* = 5.7· 10^−5^ cm^2^. In the extracellular bath surrounding the tissue sample, *u_e_* is governed by (7) with *M_i_* = 0 and *δ_e_* = 1.

#### Introducing inhomogeneities

Following [59], regional differences in the sinoatrial node tissue sample is represented by varying the conductance of the main ionic currents, *I*_Na_, *I*_CaL_, *I*_CaT_, *I*_f_, *I*_Kr_, *I*_Ks_, *I*_Kur_, *I*_NaCa_, and *I*_NaK_. For a regional variation fraction of *p* = 0.05 or *p* = 0.2, each of these parameters, *g_j_*, are set to

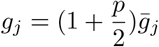

in the upper left part of the tissue sample and

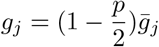

in the lower right part of the tissue sample. In the remaining parts of the tissue sample, the parameters are set to the default values, 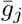.

### Numerical methods

The bidomain model equations are discretized using a finite difference discretization with Δ*x* = Δ*y* = Δ*z* = 0.02 mm and Δ*t* = 0.1 ms. The equations are solved using a standard first order operator splitting technique, splitting the non-linear dynamics of *I*_ion_ and *F* from the linear partial differential system (see, e.g., Chapter 10 of [21]). The non-linear system is solved using forward Euler with a time step of Δ*t* = 0.01 ms.

### KNM representation of a heart-on-a-chip

The KNM representation of the heart-on-a-chip with hiPSC-CMs (Figure 3) is based on [41]. The parameters, geometry and membrane dynamics (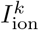 and *F_k_*) are taken directly from the baseline model representation of Chip C4 in that study. In the case of applied stimulation, we stimulate the leftmost 150 *µ*m of cells with a constant 20 ms long stimulation current of 15 *µ*A/cm^2^ at a frequency of 1 Hz.

The chip consists of a two-dimensional collection of 1347 hiPSC-CMs connected to their neighbours in the longitudinal (*x*) and transverse (*y*) directions. Each cell is assumed to have length *l_x_* = 25 *µ*m and width *d* = *l_y_* = *l_z_* = 8 *µ*m. The specific membrane capacitance is set to *C_m_* = 1 *µ*F/cm^2^ [54] and the membrane area is set to 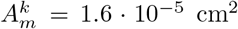 [60]. The KNM conductance parameters, 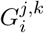 and 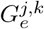, are defined by [15]

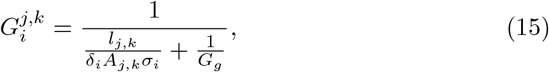

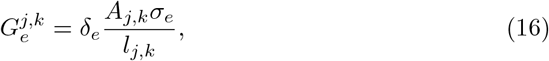

where *l_j,k_* is the distance between the centers of cells *j* and *k* and *A_j,k_* is the cross-sectional area between cells *j* an *k* (including both the intracellular and the extracellular space). These values depend on the cell geometry *l_x_, l_y_*, and *l_z_* and whether the cells are connected in the longitudinal or transverse directions (see, e.g., [40]). Furthermore, *δ_i_* = 0.8 is the intracellular volume fraction,

*δ_e_* = 1 −*δ_i_* is the extracellular volume fraction, *σ_i_* = 8.2 mS/cm [7] is the intracellular conductivity, *σ_e_* = 15 mS/cm [58] is the extracellular conductivity, and *G_g_* = 180 nS = 180· 10^−6^ mS [41] is the gap junction conductance between two neighbouring cells.

The cell collection is surrounded by an extracellular bath of approximately 200 *µ*m in each direction. In this bath, *u_e_* is governed by (10) with 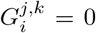and *δ_e_* = 1. A Dirichlet boundary condition of the form *u_e_* = 0 mV is applied on the outer boundary of the extracellular bath.

#### Introducing inhomogeneities

In the KNM representation of a heart-on-a-chip, we introduce cell-to-cell variation by varying the value of the conductance all membrane currents included in the model (*I*_Na_*, I*_NaL_*, I*_CaL_*, I*_to_*, I*_Kr_*, I*_Ks_*, I*_K1_*, I*_NaCa_*, I*_NaK_*, I*_pCa_*, I*_bCl_*, I*_bCa_, and *I*_f_). For each cell *k* and each conductance *j*, we draw a random value *α_k,j_* between 0 and 1. We let the conductance of current *j* for cell *k* be given by

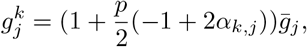

where 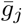is the default conductance value of current *j* and *p* is the degree of variation. For example, we use *p* = 0.1 or *p* = 0.5 for 10% or 50% cell variation, respectively.

#### Numerical methods

We solve the KNM system using the same operator splitting technique as for the bidomain model (see above). We use a global time step Δ*t* = 0.1 ms and a local time step of Δ*t* = 0.0005 ms for the forward Euler solution algorithm applied to solve the non-linear membrane dynamics system.

### KNM representation of a pancreatic islet

In the KNM representation of a pancreatic islet (Figure 4), we consider a three dimensional collection of *β*-cells with membrane dynamics (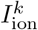 and *F_k_*) modeled by the *β*-cell model from [46]. We consider the parameterization of this membrane model corresponding to metabolically driven slow waves of electrical activity (Figure 6 of [46]). The islet consists of 1000 *β*-cells [61]. Each *β*-cell is assumed to be shaped as a sphere with diameter *d* = *l_x_* = *l_y_* = *l_z_* = 12 *µ*m [62], corresponding to a surface area of *A_m_* = *πd*^2^ ≈4.5· 10^−6^ cm^2^, and *C_m_* is set to 1 *µ*F/cm^2^ [54]. The KNM conductance parameters, 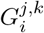 and 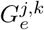, are defined by (15) and (16), and we use the same values of *σ_i_* = 8.2 mS/cm and *σ_e_* = 3 mS/cm as is applied for neurons. The extracellular volume fraction is set to *δ_e_* = 0.29 [63, 64], and the gap junction conductance between neighboring cells is set to *G_g_* = 250 pS = 0.25· 10^−6^ mS [62].

#### Introducing inhomogeneities

Cell-to-cell variation in the pancreatic islet is introduced in the same manner as for the KNM representation of a heart-on-a-chip. For the pancreatic *β*-cells, we consider variations in the three ion channel conductances *g*_Na_, *g*_CaL_, and *g*_KATP_.

#### Numerical methods

We apply the same numerical methods as for the KNM representation of a heart-on-a-chip for the KNM representation of the pancreatic islet. For the pancreatic islet, we use a global time step of Δ*t* = 1 ms and a local time step of Δ*t* = 0.1 ms for the forward Euler solution algorithm used to solve the non-linear membrane dynamics system (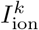 and *F_k_*).

### EMI model representation of a cerebellar purkinje neuron

The EMI model representation of a cerebellar purkinje neuron in Figures 5 and 6 is based on the cable equation representation in [7]. More specifically, the neuron is divided into different regions with specific ion channel conductances and geometries. These regions are illustrated in Figure 5a and the properties are reported in Table 1. In our EMI model representation, we have used a somewhat different geometry for the dendrites than in the original cable model representation. The rationale is that the EMI model requires a detailed three dimensional mesh (see Figure 5b) representing all membrane surfaces, and it was therefore favorable to use a dendrite geometry that was more easily generated using a simple recursive algorithm. We constructed a dendrite shaped as a tree with seven branching levels (see Figure 5a). The dendrite length and diameter used in each branching level are reported in Table 2.

**Table 2.** Lengths, *l*, (in *µ*m) and diameters, *d*, (in *µ*m) applied for each branching level in the dendrite geometry (see Figure 5).

| Level | 0 | 1 | 2 | 3 | 4 | 5 | 6 | 7 |
| --- | --- | --- | --- | --- | --- | --- | --- | --- |
| $l$ | 15 | 24 | 19.2 | 15.4 | 12.3 | 9.8 | 7.9 | 6.3 |
| $d$ | 10 | 8 | 5 | 3 | 2 | 1.5 | 1 | 1 |

Because of the change in dendrite geometry, some adjustments were made to the potassium channel conductances in the dendric regions to make the model behave like in [7] (see Table 1). In addition, in order for the largest EP spike to be generated at the axon initial segment (AIS) and not from the soma (consistent with the observations in [9]), the sodium channel conductance of the AIS was increased compared to the value used in [7] (see Table 1). Furthermore, to avoid slow convergence of calcium and buffer concentrations in response to parameter changes that could have required unfeasible simulation times for the EMI model, we have for simplicity only used the transmembrane currents from [7] and ignored the model for the intracellular calcium dynamics.

In our simulations, we have used the EMI model parameters *C_m_* = 1 *µ*F/cm^2^ [54], *σ_i_* = 8.2 mS/cm [7], and *σ_e_* = 3 mS/cm [55]. We have included approximately 100 *µ*m of extracellular space surrounding the neuron in all directions. On the outer boundary of the extracellular space, Dirichlet boundary conditions of the form *u_e_* = 0 mV are applied.

#### Introducing inhomogeneities

In the case of a uniform distribution of ion channels (left panels of Figures 5 and 6), the soma model parameters are applied on the entire membrane of the neuron. For a non-uniform distribution of ion channels, the regional differences described in Table 1 are applied in the different parts on the neuron. In the myelinated regions between the Ranvier nodes, the membrane is assumed to be insulated, and the boundary conditions **n***_e_· σ_e_*∇*u_e_* = **n***_i_· σ_i_*∇*u_i_* = 0 are applied.

#### Numerical methods

The EMI model equations are solved using the spatial and temporal operator splitting algorithm introduced in [65]. We discretize the models using the finite element method using first-order elements implemented in the MFEM finite element library [66]. The three dimensional finite element mesh is generated using gmsh [67], with mesh size ranging from about 0.5 *µ*m close to the dendrite and axon to 50 *µ*m close to the boundary of the extracellular space. We use a global time step of Δ*t* = 0.01 ms and a local time step of Δ*t* = 0.0001 ms for the forward Euler algorithm applied to solve the non-linear membrane dynamics.

